# Killer meiotic drive and dynamic evolution of the *wtf* gene family

**DOI:** 10.1101/461004

**Authors:** Michael T. Eickbush, Janet M. Young, Sarah E. Zanders

**Affiliations:** Stowers Institute for Medical Research, Kansas City, MO 64110, USA.; Division of Basic Sciences, Fred Hutchinson Cancer Research Center, Seattle, WA 98109, USA.; Department of Molecular and Integrative Physiology, University of Kansas Medical Center, Kansas City, KS 66160, USA.

**Author notes:** ^*^Correspondence to: Sarah E. Zanders, Stowers Institute for Medical Research, 1000 E 50^th^ Street, Kansas City, Missouri 64110.

## Abstract

Natural selection works best when the two alleles in a diploid organism are transmitted to offspring at equal frequencies. Despite this, selfish loci known as meiotic drivers that bias their own transmission into gametes are found throughout eukaryotes. Drive is thought to be a powerful evolutionary force, but empirical evolutionary analyses of drive systems are limited by low numbers of identified meiotic drive genes. Here, we analyze the evolution of the *wtf* gene family of *Schizosaccharomyces pombe* that contains both killer meiotic drive genes and suppressors of drive. We completed assemblies of all *wtf* genes for two S. *pombe* strains, as well as a subset of *wtf* genes from over 50 strains. We find that *wtf* copy number can vary greatly between strains, and that amino acid substitutions, expansions and contractions of DNA sequence repeats, and nonallelic gene conversion between family members all contribute to dynamic *wtf* gene evolution. This work demonstrates the power of meiotic drive to foster rapid evolution and identifies a recombination mechanism through which transposons can indirectly mobilize meiotic drivers.

## Introduction

Many genes are maintained in eukaryotic genomes by natural selection because they provide a fitness benefit to the organisms that bear them. Analyses of these genes and their molecular functions constitute the bulk of molecular biology research performed today. However, not all genetic loci provide a fitness benefit to their hosts and some can even be described as parasites. There are many types of parasitic genes, which can comprise large fractions of eukaryotic genomes and can have a substantial impact on shaping genome evolution (1).

Killer meiotic drive loci are one such class of parasites that can be particularly harmful to fitness. These selfish loci act when heterozygous to destroy the meiotic products that do not inherit them. This killing causes the heterozygote to transmit the meiotic drive locus to up to 100% of the functional meiotic products (2, 3). Killer meiotic drivers have been observed throughout eukaryotes from plants to mammals, even though their selfish behavior generally decreases overall organismal fitness (3-6). Killer meiotic drivers can directly cause infertility, and biasing allele transmission disrupts the ability of natural selection to choose the best adapted alleles at any linked loci. Genomic loci that suppress drive are therefore predicted to be favored by selection (7). Indeed, the activity of many suppressors of meiotic drive has been observed, although only four suppressor genes have been cloned (8-11).

Detecting meiotic drive and distinguishing it from other phenomena that bias allele transmission can be experimentally challenging, even in the most tractable genetic systems (1). After establishing the presence of drive loci, identifying the genes responsible often takes years. In addition, the handful of meiotic drive loci that have been cloned in different systems are not homologous to each other, so sequence analysis is generally not useful in identifying novel drivers (3-6, 12-19). These factors limit the field’s ability to efficiently analyze the possible presence or impact of meiotic drivers, especially in complex organisms with limited genetic tractability like humans.

Although meiotic drive genes generally do not share DNA sequence homology, they may share certain evolutionary signatures that could guide discovery of novel drive loci from genomic sequence data alone. For example, genetic conflict between drivers and suppressors is predicted to trigger an evolutionary arms race where both sides exhibit rapid evolution (19, 20). Similarly, evidence of analogous evolutionary arms races between viruses and host genomes has become widespread and has led to revolutionary insights in viral-host interactions (21). However, due to the paucity of cloned meiotic drivers and suppressors, studies of the evolutionary signatures of genes known to cause or suppress meiotic drive are limited (2).

The *wtf* gene family from *Schizosaccharomyces pombe* offers an exceptional opportunity to study the evolution of meiotic drive systems (22). The genomes of *S. pombe* isolates contain more than 20 *wtf* genes, some of which are known to be killer meiotic drivers (23, 24). The characterized drive genes are predicted to encode transmembrane proteins, but there are no obvious orthologs outside of S. *pombe* and the complete molecular mechanisms of drive are unknown. However, the characterized driving *wtf* genes use alternate transcripts to generate both an antidote and a poison during gametogenesis. The poison acts on all gametes, whereas the antidote remains within *wtf*+ gametes. The combined action of the poison and antidote proteins results in the preferential death of the *wtf*− gametes generated by *wtf*+/*wtf*− heterozygotes and therefore preferential transmission of *wtf*+ alleles (23, 24).

The driving *wtf* genes impose significant fertility costs on their hosts and severely limit the ability of *S. pombe* isolates to reproduce sexually (24, 25). Novel genes or genetic variants that can suppress the action of *wtf* drivers are expected to promote fitness and should be favored by natural selection (7). Consistent with this idea, a suppressor of a killer *wtf* drive gene has recently been identified. Interestingly, this suppressor, *wtf18-2*, is a member of the *wtf* family and likely evolved from a *wtf* driver (11).

In this work, we assemble and annotate the *wtf* genes from two *S. pombe* isolates and compare them to the *wtf* genes of two previously published *S. pombe* isolates (24, 26). We classify the *wtf* genes into possible functional groups based `on previously characterized genes. In addition, we greatly extend previous evolutionary analyses of the *wtf* gene family (24, 27). Consistent with their engagement in molecular arms races, we show that *wtf* genes exhibit rapid evolution. In fact, *wtf* genes are among the most rapidly-evolving genes in the *S. pombe* species group.We show that intact *wtf* gene numbers vary between isolates and that the sequences of syntenic *wtf* genes can be markedly different (<30% sequence identity), much lower than the high overall DNA sequence identity between genomes (>99%). We show that homologous recombination, repeat expansion and contraction, and positive selection for amino acid substitutions all contribute to diversification of the *wtf* gene family. This work provides a case study for the evolutionary dynamics between selfish genes and their suppressors and supports the idea that signatures of rapid evolution could guide the discovery of novel drive loci.

## Results

### Correcting *wtf* gene annotations in the *Sp* reference strain

The PomBase database provides annotated gene structures for 25 *wtf* genes, of which 3 are annotated as pseudogenes (28, 29). However, our previous analyses of the *Sp wtf4*, *Sp wtf13* and *Sp wtf18* loci revealed that the annotated splice sites were inconsistent with published long read RNA sequence data (11,23). We therefore reevaluated the remaining *Sp wtf* gene annotations using long read RNA sequence data (Supplemental Figure 1) (30). We found our predictions were consistent with the PomBase annotations for 14 *wtf* genes but different for the remaining 11 genes. Our results matched those of Hu et al. who predicted the coding sequences computationally (24). In the updated annotations, four *wtf* genes that were previously predicted to be intact (*Sp wtf6*, *wtf8*, *wtf12* and *wtf17*) are truncated by early stop codons (based on homology to other *wtf* genes). These genes join *wtf1*, *wtf2*, and *wtf3* as likely pseudogenes.

**Figure 1.**
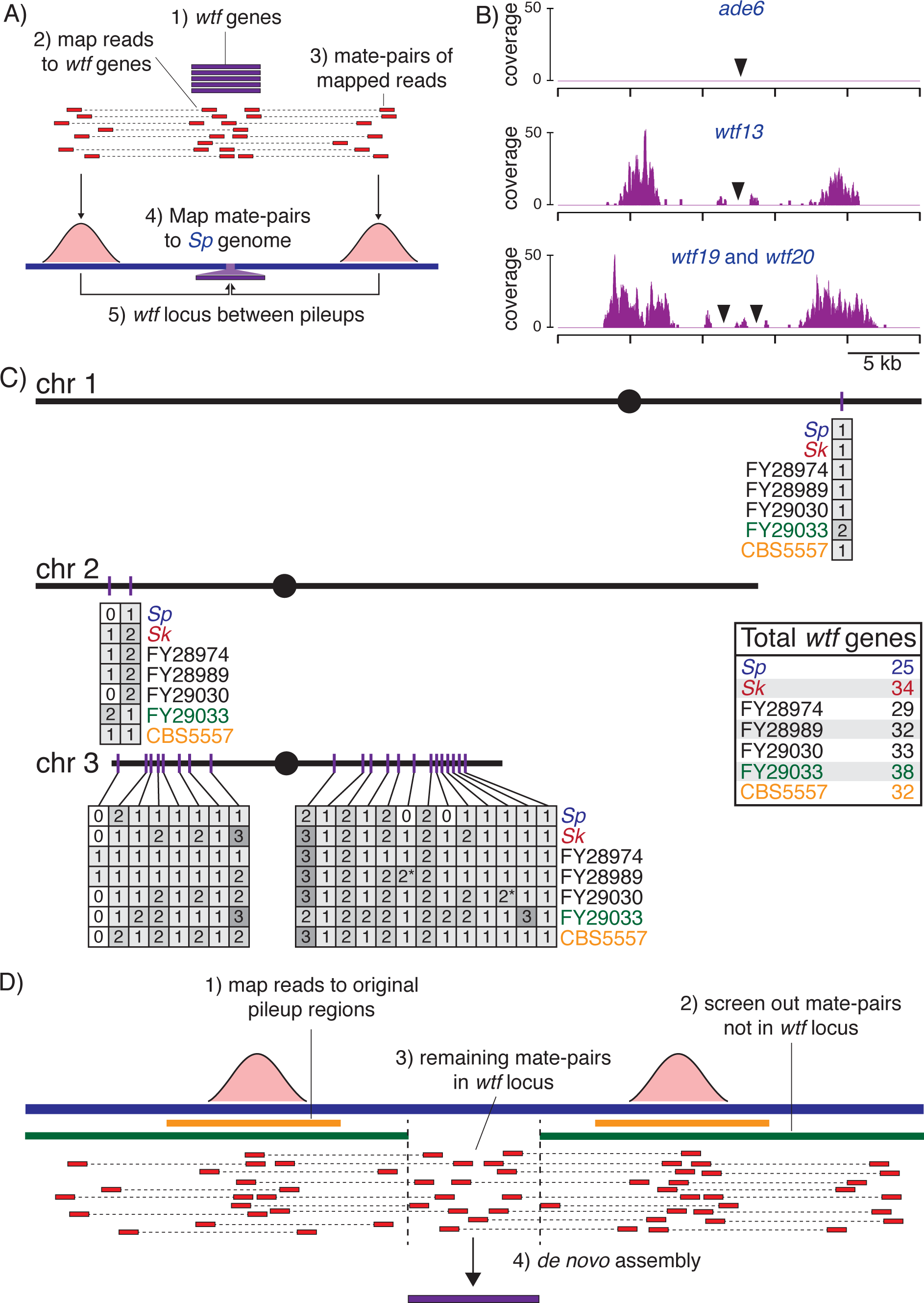
**A genomics approach identifies and assembles *wtf* gene sequences.** A) Schematic of the strategy we used to identify *wtf* gene locations. B) Examples of three *Sp* loci are shown to illustrate how read pileups (from strategy described in A) flank loci with zero, one or two *wtf* genes. In each plot, the x-axis shows relative position in the *Sp* reference genome, and the y-axis shows the number of reads mapping to each base. C) A map of *wtf* gene distribution in seven isolates of *S. pombe*. The map shows the three chromosomes of the *Sp* karyotype, although this karyotype is not shared by all isolates. The inset box indicates total *wtf* gene numbers (including pseudogenes) in each strain. The numbers for FY28974, FY28989 and FY29030 (in black) are estimates because we did not assemble all *wtf* loci in those strains. D) Schematic of the strategy we used to assemble *wtf* gene sequences in *Sk* and FY29033.

### *wtf* gene numbers vary greatly between *S. pombe* species group isolates

The molecular arms race model predicts that genes in conflict, such as meiotic drivers and their suppressors, will evolve rapidly in order to outcompete one another (20). Gene duplication is a commonly used evolutionary strategy to facilitate rapid diversification and has been observed in the context of virus-host arms races (21). The large number of *wtf* loci in the reference *S. pombe* genome assembly (25 genes, including pseudogenes) is consistent with a similar scenario occurring within the *wtf* family. In addition, previous limited analyses revealed differing numbers of *wtf* genes between different *S. pombe* group isolates (23, 24). To more globally test the possibility that recent *wtf* gene duplications or deletions have occurred in the *S. pombe* group, we first determined whether *wtf* gene numbers are dynamic between strains.

In addition to the reference *S. pombe* strain (972, isolated in France in 1921), over 150 genomes of *S. pombe* isolates have been sequenced using paired-end 100 base pair Illumina reads with standard insert sizes (~300 base pairs) (31-33). In addition, the genome of the CBS5557 strain (collected in Spain, reported 1964) was also analyzed using long-read PacBio sequencing (24). Due to the repetitive nature of the *wtf* genes and the fact that they are often flanked by repetitive Tf transposons or Tf long terminal repeats (LTRs), the sequences of *wtf* loci could be reliably determined in CBS5557 using long reads, but not in genomes where only short reads were available. To overcome this challenge, we sequenced five *S. pombe* isolates and the reference strain using ‘mate-pair’ libraries to capture pairs of 150 bp reads separated by 5-8 kb in the genome (Supplemental Figure 2). We obtained >80X coverage of each genome. With this large insert sequencing approach, the distance between mate pair reads is large enough that when one read of the pair falls within a repetitive *wtf*, the mate often falls in unique genomic sequence (Figure 1A). This allowed us to match *wtf* reads with their cognate genomic locus, even for *wtf* genes that share very high sequence identity. We sequenced derivatives of the *S. pombe* reference strain (which we will abbreviate as *Sp*), FY28974 (collected in Brazil in 1996), FY28989 (collected in Africa in 1921), FY29030 (collected in Indonesia in 1949), FY29033 (collected in Indonesia in 1923), and *Schizosaccharomyces kambucha* (abbreviated *Sk*, isolated in the USA, reported in 2002) (26, 33, 34). *Sk* was historically given a different species name because it is reproductively isolated from *Sp*, but it is no more diverged from *Sp* than other isolates of the *S. pombe* group (25, 34, 35). Like all isolates classified as *S. pombe*, all strains analyzed in this work are all very closely related: they are estimated to have diverged from each other within the last ~2,300 years and share on average >99% DNA sequence identity (32).

**Figure 2.**
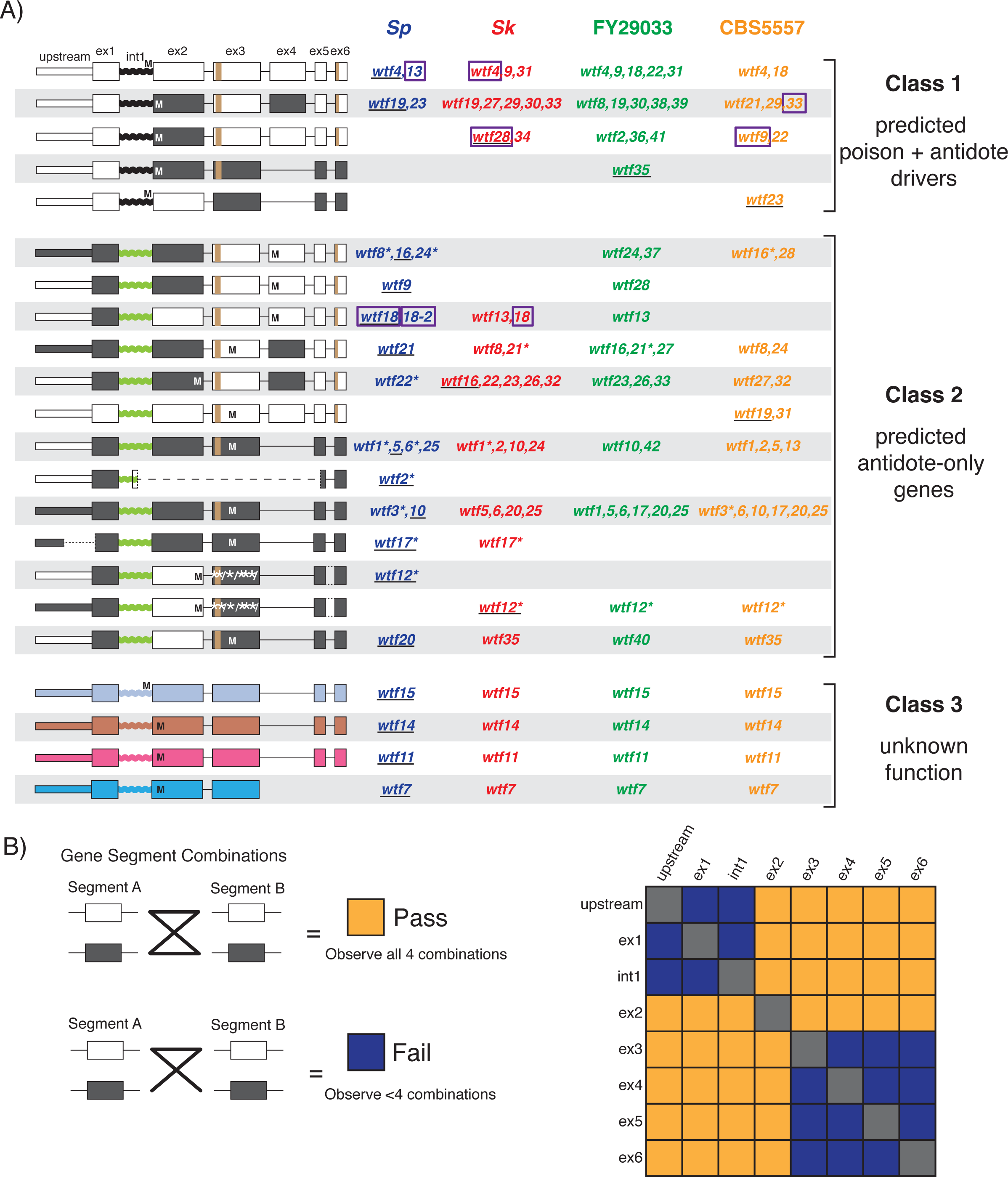
**Classification of *wtf* genes based on sequence, and evidence of nonallelic gene conversion.** A) Although they are quite diverged from each other, *wtf7*, *wtf11*, *wtf14* and *wtf15* were placed in a shared class because their sequences are unlike any functionally characterized genes. For the remaining genes, individual gene segments (each exon, intron 1 and the upstream regions) from all genes were aligned and classified based on the major clades in maximum likelihood trees (see text for details). Each segment’s clades were color-coded for display purposes (i.e. black/white, black/green coding), and genes were grouped based on gene segment patterns. On the left, we display cartoons of gene structures for each group, with ‘M’ indicating in-frame start codons, ‘/’ indicating frameshift mutations, and ‘^*^’ indicating in-frame stop codons. The repeat regions found in exons 3 and 6 are shown in brown. The names of genes in each class are listed on the right, with the gene illustrated in the cartoon underlined, and pseudogenes denoted with asterisks after gene names. The predicted function of each gene class is shown on the far right. Genes with experimentally verified phenotypes have their names outlined with purple boxes. B) Pairwise four-gamete test for recombination (gene conversion) between all pairs of *wtf* gene segments for the genes in Classes 1 and 2. Orange boxes indicate that recombination likely occurred because all four segment combinations were observed. Purple boxes indicate that not all segment combinations were observed.

To identify genomic loci in each strain that harbor *wtf* genes, we first used our sequence data to select all read pairs in which one of the reads aligned to one or more of the 25 *wtf* genes in the reference genome (abbreviated as *Sp* here). We then isolated mates of those *wtf* reads, aligned them to the *Sp* reference genome, and visually analyzed regions where multiple *wtf* mate reads mapped (‘pileups’). This yielded a map in which each *wtf* locus is flanked by pileups of mate reads that map uniquely in the genome (Figure 1A and 1B). To verify this approach, we applied it to the *Sp* data and accurately detected all *wtf* locations. We further observed that *Sp* loci containing a single *wtf* gene were typically flanked by ~ 2.2 kb wide pileups, slightly wider than the typical genomic width of a *wtf* gene (average 1.2 kb). *Sp* loci encoding two *wtf* genes were flanked by wider (~4.4 kb) pileups (Figure 1B, Supplemental Figure 3). These data suggested we could use the presence and width of such pileups to identify *wtf* loci genome-wide.

**Figure 3.**
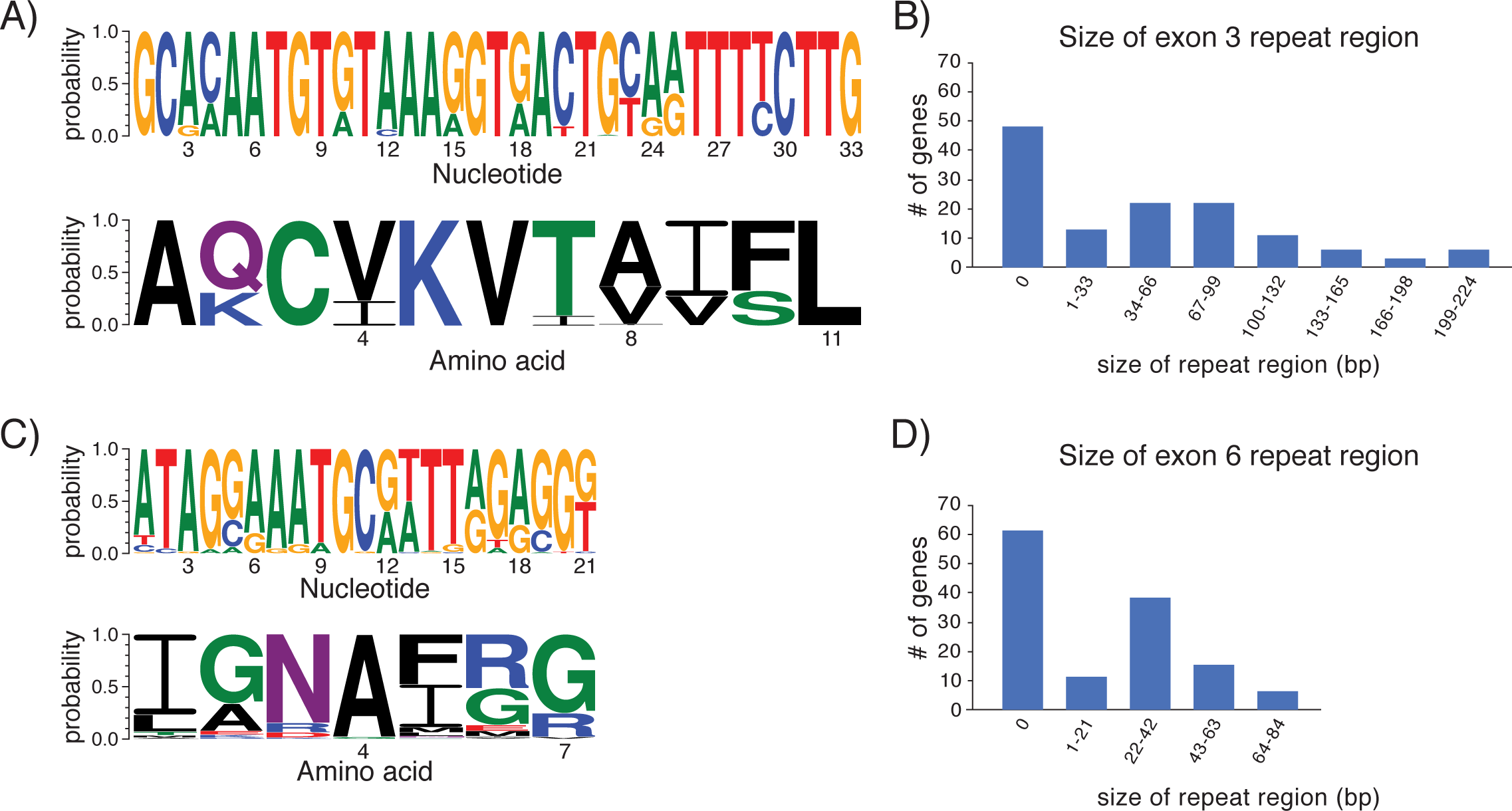
**Expansion and contraction of repeat sequences contributes to rapid *wtf* gene evolution.** A) DNA (top) and amino acid (bottom) sequence logos representing the repeat region found in exon 3. B) The distribution of exon 3 repeat region size across all assembled *wtf* genes. The sizes are presented in base pairs instead of repeat units because the initial and terminal repeats are not always full length. C) DNA (top) and amino acid (bottom) sequence logos representing the exon 6 repeat region found in many *wtf* genes. D) The distribution of exon 6 repeat sizes in all assembled *wtf* genes.

We then used this approach to identify *wtf* loci in each of the five strains we sequenced and to estimate how many *wtf* genes each locus contains (Fig 1C). In *Sk* and FY29033, these estimates were confirmed (and in a few cases corrected) by assembly of the *wtf* loci from the mate-pair reads (see below) and by Sanger sequencing of some *Sk* loci (*wtf7*, *wtf9*, *wtf13*, *wtf14+wtf15*, *wtf17+18*, *wtf33*, *wtf19+20*, *wtf23*, *wtf27*, and *wtf35*). Unlike in *Sp*, in each of the other strains we found a few loci flanked by even wider pileups (up to~7.5 kb), suggesting that these loci each contain three *wtf* genes (Supplemental Figure 3). This inference was confirmed by assembling 4 such sites.

At most of the loci we detected, we observed a symmetrical pair of pileups that were ~2.2 or ~4.4 kb wide that clearly suggested one or two *wtf* genes within the locus. Some loci, including most of those with three *wtf* genes, showed more complicated or misleading patterns (examples are shown in Supplemental Figure 4). Assembly of these regions in *Sk* and FY29033 revealed these inconsistencies were due to transposon insertions near the *wtf* loci that were not present in the reference genome. For example, a transposon insertion meant that the two genes at the *wtf2* locus in *Sk* showed a pileup pattern typical of a one gene locus, and an additional transposon meant that the three genes at the *wtf10* locus in *Sk* showed a pileup pattern typical of a two gene locus (Supplemental Figure 4). The errors in our FY29033 and *Sk* gene number predictions (at 3 out of 46 sites) were detected during the assembly of those loci to obtain *wtf* gene sequences (below). As we did not assemble all *wtf* loci in FY28974, FY28989 and FY29030, there could be similar uncorrected underestimates of *wtf* gene numbers in those strains. In addition, our method would be unable to detect more than three tandem *wtf* genes because the locus size exceeds the insert size between our mate-pair reads (Supplemental Figure 2). Although we did not observe loci with more than three tandem *wtf* genes in genomes with assembled *wtf* loci, this limitation could also lead to an underestimate of *wtf* gene numbers in the genomes where we did not perform *de novo* assemblies.

**Figure 4.**
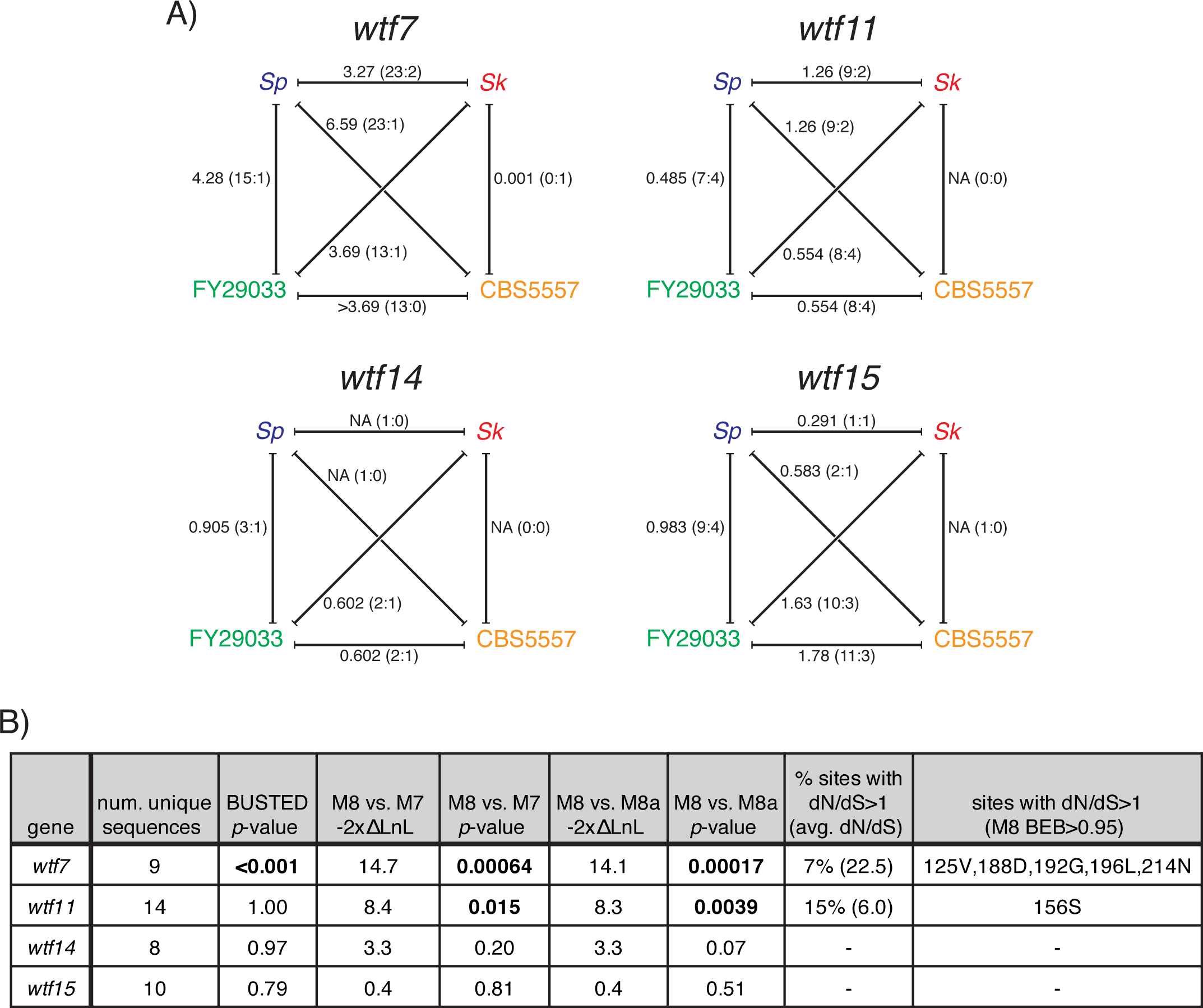
**Analysis of selective forces acting on *wtf7*, *wtf11*, *wtf14* and *wtf15* genes.** A) dN/dS analyses for all pairwise combinations between orthologous genes. The dN/dS ratio is shown, with the numbers of nonsynonymous and synonymous changes shown as a ratio in brackets. Alignments highlighting the variant codons are shown in Supplemental Figures 17-20. B) BUSTED and PAML analyses of alleles from 57 strains. P-values supporting positive selection are highlighted in bold (see text for details).

Our mate-pair pileup approach could also miss additional *wtf* gene copies if they were found within larger recently duplicated regions of the genome. To look for such *wtf* genes, we aligned all sequence reads for each strain to the *Sp* reference genome and looked for regions containing *wtf* loci where sequencing coverage was roughly twice as high as the rest of the genome. We found two duplicated regions that include a *wtf* gene (Supplemental Figure 5). In FY29030, there is a ~14 kb duplication of the *wtf23* region (between chromosome 3 reference genome positions 2,145,417 and 2,159,329). In FY28989, there is a ~95 kb duplication of the *wtf33* region between positions 1,838,980 and 1,933,773 on chromosome 3 (Supplemental Figure 5). These duplications both appear to be very young, as we do not detect increased sequence variation in thosex regions compared to the flanking sequence. We therefore conclude that FY29030 contains two nearly identical copies of *wtf23* and FY28989 contains two nearly identical copies of *wtf33* (indicated with asterisks in Figure 1C).

After identifying all the genomic loci encoding *wtf* genes in the five strains, we combined our data with the previously identified *wtf* landscapes in CBS5557 and *Sp* (24, 29). Altogether, we found that the total number of *wtf* genes (including pseudogenes) varied greatly between strains, ranging from 25 in *Sp* to 38 in FY29033 (Figure 1C). Each locus can contain between zero and three *wtf* genes. Overall, the locations of *wtf* genes were quite similar between isolates: we found only four *wtf* loci that were not shared among all strains. Most of the variation in *wtf* number between strains can be explained by expansion/contraction of *wtf* gene numbers within each locus (Figure 1C), although without a clear outgroup strain it is unclear what the relative contributions of duplications and deletions are. Given that all strains encode at least one *wtf* gene at 20 shared loci, it is likely that the ancestral genome of these strains contained at least 20 *wtf* genes.

### Assembling *wtf* genes from *Sk* and *FY29033* yields many unique gene sequences

Within *Sp*, there is extensive sequence diversity amongst the *wtf* genes. Some, like *Sp wtf4* and *Sp wtf13*, are very similar (>90% amino acid identity), whereas others, like *Sp wtf4* and *Sp wtf7*, are not (<30% amino acid identity). We wanted to know if the gene repertoire of *Sp* reflects the full range of *wtf* diversity, or if it instead represents a limited sample. To test this, we used our sequencing data to assemble all *wtf* genes from two additional genomes, *Sk* and FY29033. We assembled each *wtf* locus separately, first selecting all read pairs in which one of the reads aligned to a unique wtf-flanking region (i.e. the pileup regions discussed above, Figures 1A, 1B and 1D). We then assembled those read pairs to generate a contig with the *wtf* gene(s) in the center (Figure 1D).

To validate this approach, we also used it to assemble *Sk wtf* genes we had previously Sanger sequenced (23). We found that our assembly matched the Sanger sequencing in most cases, but differed at the *Sk wtf2* locus. Further analyses revealed that our new assembly of the *Sk wtf2* region was correct and the Sanger sequencing missed a Tf transposon and a second *wtf* gene (*wtf34*) in the region, likely due to template switching during PCR amplification. These results suggest our assembly approach is robust.

We then predicted *wtf* coding sequences based on possible open reading frames and homology to annotated *Sp wtf* genes. Our analyses (discussed below) found additional *wtf* gene variation not represented in the *wtf* genes found in *Sp* or CBS5557.

### Naming *wtf* genes

There are currently three reported phenotypic classes of intact *wtf* genes: killer meiotic drivers, suppressors of drive, and one essential gene (*Sp wtf21*) (11, 23, 24, 36). It is unknown if there are other phenotypic classes of *wtf* genes, but it would not be surprising given their vast diversity. To facilitate answering this question and to guide future phenotypic classification of *wtf* genes, we assigned gene names to each *wtf* gene from *Sk*, FY29033, and CBS5557.

For *Sp*, we used existing gene names, and for each other genome, we named genes according to their genomic synteny by comparison with *Sp*. We use *Sk* as an example to explain our naming scheme. At the loci where both *Sk* and *Sp* have one *wtf* gene, we gave the *Sk* gene the same number as *Sp* (e.g., *Sk wtf1*), regardless of sequence identity. For loci where *Sp* has one gene and *Sk* has two genes (e.g., at the *Sp wtf8* locus), we gave the same gene number to the *Sk* gene that was most similar to the *Sp* gene and gave the remaining *Sk wtf* genes increasing numbers (26-35) depending on their order in the *Sk* genome. We followed the same convention for naming the FY29033 and CBS5557 *wtf* genes to facilitate comparisons between strains (the genes of CBS5557 were already named by Hu et al. (24) as cw1-cw29; we provide name translations in Supplemental Table 1). Supplemental Figure 6 shows *wtf* gene names and locations in the four strains.

### Pervasive nonallelic gene conversion between *wtf* genes

To examine *wtf* gene evolution, we aligned their coding sequences and generated a maximum likelihood phylogenetic tree. Naively, we expected that sets of genes from the four sequenced strains that are found in syntenic loci would group together in well-supported clades on the tree. However, syntenic genes grouped with one another in only a few clades of the tree. The *wtf7*, *wtfl1*, *wtf14*, and *wtf15* genes each form well-supported clades that do not include genes from other loci (bootstrap values >95%; Supplemental data and Supplemental Figure 7). Each of these genes is quite distinct from other *wtf* genes (separated by long branches). The alleles of the *wtf12* and *wtf17* genes also form well-supported clades (>80% support), albeit less diverged from their nearest neighbors (Supplemental Figure 7). These genes however, appear to be losing function in at least some strains: shared inactivating mutations in the *wtf12* gene in all four strains indicate that it pseudogenized prior to the divergence of the strains, and the *Sp* and *Sk* sequences of *wtf17* also appear pseudogenized.

Despite clear synteny and a very short time (~2,300 years) since these yeast isolates shared a common ancestor (32), none of the remaining syntenic *wtf* gene sets form well-supported clades that exclude *wtf* genes from other loci. Furthermore, there are clear examples of well-supported clades containing genes from different loci. For example, one well-supported clade includes the following genes: *Sk wtf29* and *wtf30*; *Sp wtf19* and *wtf23*; FY29033 *wtf8*, *wtf30* and *wtf38*; and CBS5557 *wtf29* (highlighted in Supplemental Figure 7). Finally, the tree contains two well-supported terminal nodes in which gene pairs at distinct loci from the same isolate (*Sp wtf19* and *wtf23* as well as *FY29033 wtf1* and *wtf35*) form a clade, while syntenic genes from other isolates are in distinct clades. These observations are consistent with gene conversion within the *wtf* gene family.

To analyze whether entire *wtf* coding sequences might be over-writing one another by gene conversion, or whether only portions of the genes are involved, we performed GARD (Genetic Algorithm for Recombination Detection) analysis on our coding sequence alignment to test for recombination between *wtf* genes (Supplemental Figure 8) (37). This algorithm tests the hypothesis that the same phylogenetic tree represents the entire alignment or if different trees best represent different segments due to recombination. GARD analysis found that the hypothesis of multiple segments with different trees was >100 times more likely than the hypothesis of a single tree. In addition, GARD identified three likely segments (p<0.01, Supplemental Figure 8). Together, our observations are consistent with widespread nonallelic gene conversion between members of the *wtf* gene family excluding *wtf7*, *wtf11*, *wtf14* and *wtf15*. Such gene conversion obscures the evolutionary history of the *wtf* gene family and means that functional inferences can often not be made across strains based on shared synteny. This work confirms and expands observations made by Hu et al. who previously described gene conversion among *Sp* and CBS5557 *wtf* genes (24).

To further explore within-wtf gene conversion, we compared the genes (excluding *wtf7*, *wtf11*, *wtf14* and *wtf15*) in segments. Most *wtf* genes have either five or six exons. For ease of comparison, we named the exons 1-6 based on the longest *wtf* genes (Figure 2). The five-exon genes are missing ‘exon 4’ but the remaining exons can be aligned to those of the six-exon genes (Figure 2). After excluding repetitive regions in exons 3 and 6 (discussed below), we generated alignments and trees (Supplemental figures 9-16, Supplemental Data) for each exon separately. We also generated alignments and trees for a conserved region (133-289 base pairs) upstream of the start codon and for intron 1, regions that in intact *wtf* drive genes presumably contain the promoters for the antidote and poison transcripts, respectively (23, 27). The division between segments along intron/exon boundaries was arbitrary: there is no reason that gene conversion should show breakpoints at these boundaries.

Strikingly, trees made from different gene segments do not show the same topology as one another (Supplemental figures 6-19, Supplemental Data). Although the short length of each segment means that bootstrap support values are generally low throughout the trees, each tree shows a broad subdivision between two main clades of *wtf* genes. For all but the shortest segment (exon 5), these two main clades are separated by a node with high bootstrap support. However, for different gene segments, the two main clades group different subsets of genes together. For example, while for exon 3, *Sp wtf9* and *Sk wtf9* group together very closely in the tree and share 96% nucleotide identity, in contrast for exon 2 they are in different main clades (separated by a well-supported node) and show very remote homology (Supplemental figures 10 and 11). One possible explanation for this pattern is that their high similarity in exon 3 reflects their original syntenic relationship, but that relationship has been obscured in exon 2 by gene conversion from another *wtf* gene overwriting sequence in one or both of the strains.

We used the broad clade divisions defined by the trees for each segment to generate a cartoon representation of this ‘patchwork’ evolutionary history. In the cartoon, each color represents one of the two well supported clades for each gene segment (Figure 2A). We used the color coding to guide grouping the full length *wtf* genes as shown (Figure 2A). We then carried out four-gamete tests to look for evidence of gene conversion between the gene segments (38). Briefly, we considered each of the two major clades for each segment as alternate alleles. We then did pairwise comparisons of all gene segments to assay how many of the four possible allele (clade) combinations were observed. The four-gamete test is positive when all four combinations are present; while a simple accumulation of individual sequence changes could explain up to three combinations, the fourth combination can only be explained by recombination (Figure 2B). We found that 19/28 comparisons yielded a positive four-gamete test. While we cannot reconstruct the full history and exact boundaries of gene conversion among *wtf* genes in each strain, it is clear that the gene family has experienced rampant sequence exchange that could have facilitated rapid functional divergence of the gene family by bringing together new combinations of sequence variants.

### DNA double strand break hotspots are enriched near *wtf* genes

The high level of nonallelic gene conversion among *wtf* genes is surprising because nonallelic homologous recombination (also known as ectopic recombination) is thought to be generally suppressed (39). This suppression is important because recombination events between nonallelic loci can result in genetic exchanges (crossovers) that cause deleterious chromosome rearrangements (39). The gene conversion among *wtf* genes we observe could be caused by increased frequency of nonallelic homologous recombination amongst these genes, or due to selection favoring the products of gene conversion events. The two explanations are not mutually exclusive and both could contribute. The latter idea is difficult to test, so we focused on the first idea. Gene conversion results from the repair of DNA double strand breaks (DSBs). The initiating DSB could happen near or within the gene converted locus itself, or the break could happen in a different (donor) site that shares homology with the gene converted locus (e.g., another similar *wtf* gene) (39). DSBs arise at low frequencies in vegetative cells, but are dramatically induced (~58 breaks per cell in *Sp*) during meiosis (40). Due to their greater numbers and the fact that they have been mapped, we focused our analyses on meiotic DSBs.

Meiotic DSBs do not form randomly and are instead enriched in regions called ‘hotspots.’ *Sp* has 602 DSB hotspots that are generally conserved between *Sp* and *Sk*, so it is reasonable to assume the *Sp* hotspot map represents the *S. pombe* group (25, 40). The *wtf* genes could have elevated gene conversion frequencies if all or a subset of them are near DSB hotspots. A factor known as the ‘gene conversion tract length’ would affect how near to a break *wtf* genes must be in order to be involved in gene conversion events as either donors or recipients. This tract length specifies the amount of DNA that may be incorporated in the DSB repair event and potentially involved in gene conversion. The gene conversion tract length has only been coarsely measured in *Sp* for allelic meiotic recombination at one locus (*ade6*). The observed gene conversion tract lengths were generally less than 1 kb and occasionally >2 kb (41). It is unknown if gene conversion varies by locus, and whether tract length is different for allelic repair than for nonallelic recombination. Given this high level of uncertainty, we designated hotspots within 2.5 kb of a *wtf* gene as potential sources of initiating gene conversion events.

We looked for an association between the 602 previously defined *Sp* DSB hotspots and *wtf* loci by calculating the distance between each end of the *wtf* coding sequences and the nearest DSB hotspot (40). There was no DSB hotspot 5’ to the first *wtf* gene on chromosome 3, so we only considered the hotspot 3’ of this coding sequence yielding 47 data points (2 ends of each of the 24 loci containing *wtf* genes minus 1). We did the same comparison for all annotated coding sequences (29). We found that DSB hotspots were significantly enriched within 2.5 kb of *wtf* loci as compared to all coding sequences. This enrichment was also significant if we only considered hotspots within 1 kb (Table 1 G-test p<0.01). Overall, we found that 14 of the 24 *wtf* loci are within 2.5 kb of one or more hotspots.

**Table 1.**
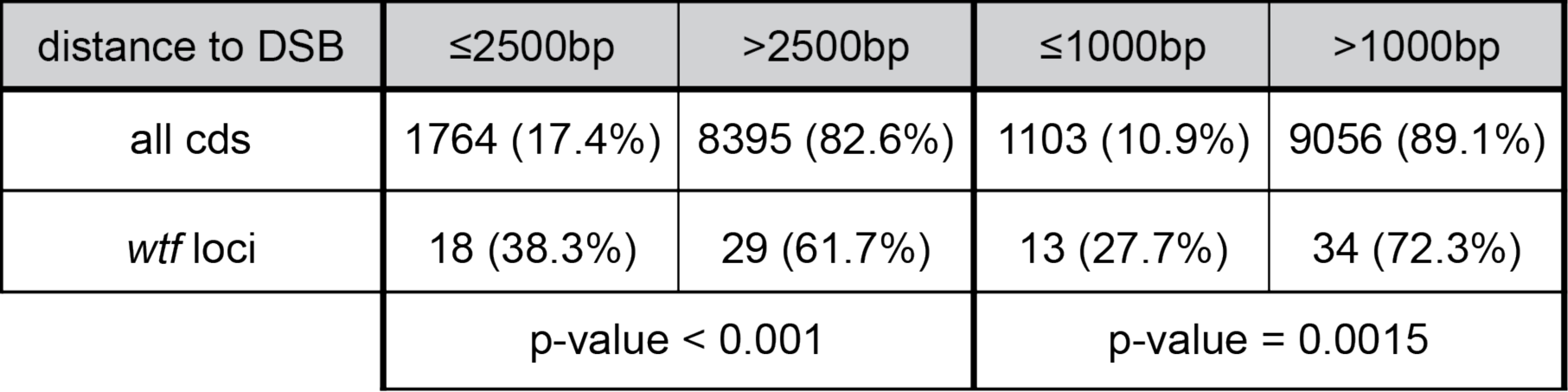
DSB hotspots are enriched near *wtf* loci

| distance to DSB | ≤2500bp | >2500bp | ≤1000bp | >1000bp |
| --- | --- | --- | --- | --- |
| all cds | 1764 (17.4%) | 8395 (82.6%) | 1103 (10.9%) | 9056 (89.1%) |
| *wtf* loci | 18 (38.3%) | 29 (61.7%) | 13 (27.7%) | 34 (72.3%) |
|  | p-value < 0.001 |  | p-value = 0.0015 |  |

These analyses suggest that close proximity of some genes to DSB hotspots likely contributes to the high levels of recombination within the *wtf* gene family. Interestingly, we observed no chromosome rearrangements with breakpoints in *wtf* genes in the 4 strains with assembled *wtf* genes despite the hotspots and evidence of gene conversion. This suggests that nonallelic homologous recombination events are either preferentially repaired as gene conversions, as opposed to crossovers, or that strains resulting from such crossovers have been removed by selection because they often generate chromosomes missing essential genes and/or with inviable duplications.

### High diversity of intragenic repeats in *wtf* proteins

Insertions and deletions within genes can be an additional source of evolutionary novelty that can result from errors during DNA replication or from recombination (42). We looked for evidence of such changes within *wtf* genes and found two repetitive regions that have frequently expanded and contracted during *wtf* evolution. The first of these is a region containing a well-conserved 33 base pair repeat sequence near the beginning of exon 3 in most *wtf* genes (Figure 3A). Not all of the repeat units are complete. The first repeat is routinely truncated to 21 nucleotides, while the last repeat is truncated to between 14 and 26 nucleotides. The *wtf* genes have between 0 and 224 bp of sequence derived from this repeat (Figure 3B). A second dynamic repeat region occurs at the start of exon 6 in most genes in Classes 1 and 2 (Figure 3C). This 21 base pair repeat unit is less conserved and not all repeat units are complete. This repeat comprises between 0 and 84 bp of sequence in *wtf* genes (Figure 3D). Both repeat regions appear unstable in that *wtf* alleles that are otherwise similar can vary in the number of repeat units. For example, the *Sp* and *Sk* alleles of *wtf4* are 93% identical outside of the repeats, but have different copy numbers of both repeat segments. These changes may be functionally important because the repeats often overlap predicted transmembrane domains. The function of these repeats is currently unknown, but the number of repeats found in exon 6 can be important for conferring specificity between poison and antidote proteins (11).

### Positive selection implicates *wtf7* and *wtf11* in genetic conflict

It is clear that gene duplication, deletion, gene conversion and changes in repeat units have all acted to generate extensive diversity in the *wtf* gene family. We also wondered whether individual amino acid changes have also played a role in increasing *wtf* diversity. We therefore analyzed the relationship between the number of nonsynonymous changes per nonsynonymous site (dN) and the number of synonymous changes per synonymous site (dS). In these analyses, dN/dS ratios near 1 are consistent with neutral protein evolution, meaning that amino acid changes in the gene are neither selected for or against. Alternatively, ratios that deviate significantly from 1 are expected when a gene is evolving under selection. Ratios less than 1 are consistent with purifying selection, meaning that novel protein variants are selected against. For example, histone genes show very low dN/dS ratios, because novel variants in histones are generally deleterious and are thus removed by purifying selection. Ratios greater than 1 are consistent with positive (diversifying) selection, meaning that novel protein variants have a selective advantage. Genes involved in genetic conflict often show signatures of positive selection (20, 21).

Unfortunately, our ability to perform dN/dS analysis in the *wtf* family is limited because the widespread nonallelic gene conversion (described above) we observe among *wtf* genes can seriously confound dN/dS analyses. However, we performed dN/dS analyses on the four *wtf* genes that appear to have escaped the effects of gene conversion: *wtf7*, *wtf11*, *wtf14* and *wtf15* (Figure 4A, Supplemental Figures 17-20). We found that some orthologous gene pairs were identical or nearly identical between isolates. For example, we found very few mutations distinguishing the *wtf14* orthologs. For *wtf11* and *wtf15*, pairwise comparisons between the orthologs revealed dN/dS values less than 1 for some pairs and dN/dS greater than 1 for others, but the overall number of changes was low, limiting the power of these analyses. For *wtf7*, however, more changes have accumulated, and there were strong signatures of positive selection: all pairwise comparisons with more than one mutation had dN/dS greater than 3 (Figure 4A).

Overall, our dN/dS analyses were limited by the small number of isolates we assayed and in some cases the low number of codon changes. These small numbers make actual deviations of dN/dS from the neutral expectation difficult to distinguish from random fluctuations. To overcome this limitation, we assembled the sequences of *wtf7*, *wtf11*, *wtf14* and *wtf15* from 54 additional *S. pombe* isolates using published 100 base pair paired-end read data (33). This was possible due to the large divergence between each of these genes and all other *wtf* genes. In many cases, the sequences of orthologous genes were identical between the strains. After removing redundant sequences, we were left with 9 alleles of *wtf7*, 14 alleles of *wtf11*, 8 alleles of *wtf14* and 10 alleles of *wtf15*. We aligned these sequences and screened each alignment for evidence of gene conversion using the GARD algorithm (37), but did not observe signatures of gene conversion.

We then tested each alignment for evidence of positive selection (Figure 4B). First, we used the codeml algorithm from the PAML (phylogenetic analyses by maximum likelihood) package (43) to test for positive selection on a subset of codons in each gene. We found statistical support for positive selection in both *wtf7* and *wtf11*, but not *wtf14* or *wtf15* (Figure 4B). We next used the BUSTED (Branch-site Unrestricted Statistical Test for Episodic Diversification) algorithm from the HyPhy suite which tests for evidence of selection on at least a subset of codons in at least a subset of the included sequences. BUSTED found support for positive selection on a subset of codons in *wtf7*, but not in *wtf11*, *wtf14* or *wtf15* (Figure 4B) (44). Overall these analyses support the hypothesis that *wtf7* and *wtf11* are engaged in genetic conflicts.

### Some *wtf* genes show characteristics of poison-antidote systems, whereas others may encode antidote-only suppressors

In addition to facilitating visualization of gene conversion, we grouped the *wtf* genes into the three major classes shown in Figure 2 to guide future functional analyses. Briefly, we dubbed the genes that contain in-frame start codons just upstream or near the beginning of exon 2 ‘Class 1’ genes. These exon 2 ATG codons encode the start of Wtf poison protein isoforms and are shared by all of the previously known drivers (Figure 2) (11,23,24). In addition, we used published long read RNA sequences to confirm that all the *Sp* Class 1 genes have an alternate transcriptional start site within intron 1 (Supplemental Figure 1) that could encode poison transcript isoforms (30). We therefore predict that Class 1 genes are intact meiotic drivers in which transcripts that include all exons encode antidote proteins, and transcripts which exclude exon 1 encode poison proteins.

Most other genes lack both a transcriptional isoform that excludes exon 1 and an in-frame ATG near the start of exon 2: we classify these as Class 2 genes. Due to similarity between the Class 2 genes and the antidote proteins produced by known drivers, we predict these genes are suppressors of *wtf* drive genes and lack the poison isoform (22). Indeed, Class 2 contains the only known *wtf* drive suppressor, *Sp wtf18-2* (11). Consistent with the predicted lack of poison isoform, the *Sk wtf5* and *wtf6* genes do not cause drive in *Sp* (23). Notably, we found no *wtf* genes that lack exon 1 that would encode only poison isoforms: it would have been very surprising to find such genes as we predict they would encode ‘suicide’ alleles unless they were very closely linked to a completely effective suppressor.

Class 3 consists of the remaining genes: *wtf7*, *wtf11*, *wtf14* and *wtf15*. These genes are diverse and are grouped together only because they all have unknown functions. These genes do have an in-frame start codon near the start of exon 2, like known drivers. However, long read RNA sequencing data showed no evidence of alternate transcripts for these genes beginning in intron 1 (Supplemental Figure 1), so we cannot make a clear prediction about whether they actually encode poison isoforms (30). Furthermore, their increased sequence divergence from the rest of the *wtf* family could suggest divergent functions.

## Discussion

Our study extends previous evolutionary analyses to demonstrate extremely dynamic evolution of the *wtf* gene family in multiple lineages of *S. pombe* (24). Although the genomes of different isolates of the *S. pombe* group are nearly identical (>99.5% DNA sequence identity) (32), the number of *wtf* genes (including pseudogenes) found in the different isolates we studied is variable and the sequences of syntenic genes can be very diverged. This rapid evolution scenario is consistent with molecular arms race models that predict rapid evolution of meiotic drivers and their suppressors (20). It also supports the idea that rapid evolution could be a hallmark of these genes that could be used, along with other features like germline expression and lineage restriction, to facilitate their discovery.

### Model for *wtf* family expansion on chromosome 3

As noted by Bowen et al., the introns found in all *wtf* genes argue against gene family expansion by retrotransposition (27). These authors also suggested that some *wtf* genes co-duplicated with their associated LTRs. In other words, *wtf* genes took advantage of the ubiquity of distributed transposon sequences to spread within the genome via nonallelic gene conversion to preexisting LTRs, a process known as segmental duplication (45). As most *wtf* loci contain at least one *wtf* gene in the majority of the seven isolates analyzed here, we propose that the segmental duplications of *wtf* genes largely occurred prior to the divergence of these isolates and perhaps the *S. pombe* group.

The exploitation of distributed transposon sequences to facilitate the spread of meiotic drivers may not be specific to *wtf* genes. Transposon sequences are also found near *Spok* genes, a different family of single-gene killer meiotic drivers in the fungus *Podospora anserina. Spok* genes are found in as many as 11 copies per genome in some species of fungi (9). Although it is unknown whether *Spok* genes are associated with transposons in other species, segmental duplication to preexisting transposon sequences may have also facilitated growth of the *Spok* gene family.

In addition to segmental duplication, tandem duplications (and deletions) also appear to have contributed to the expansion (and contraction) of the *wtf* gene family. Nonallelic recombination and slippage during DNA replication could be contributing to duplications and deletions. These events appear to have continued after the divergence of the strains analyzed here because the number of *wtf* genes at any given locus varies (Figure 1C). For example, Hu et al. found that *wtf27*, *wtf33*, and *wtf35* gene were all apparently lost in the *Sp* isolate due to recombination between two LTRs in the same orientation that flanked the genes (24).

Interestingly, like in the reference genome (*Sp*), the *wtf* genes in all the strains assayed are highly enriched on what is chromosome 3 in *Sp*. Bowen et al. proposed that this enrichment in *Sp* could reflect a different evolutionary origin for chromosome 3, suggesting that it was introgressed from a diverged strain with many *wtf* genes throughout the genome (27). If this is true, such an introgression event must have preceded the divergence of the strains analyzed here (Figure 1). We have proposed an alternative hypothesis, that the segmental duplication events spreading *wtf* genes occur genome-wide, but that the duplicates on chromosome 3 are preferentially maintained, because *S. pombe* can tolerate aneuploidy of only chromosome 3 and not the other chromosomes (22). This could be important because when two or more distinct *wtf* drivers compete (i.e. they are linked on opposite haplotypes), nearly all haploid gametes are expected to be destroyed. This was, in fact, observed when CSBS5557 *wtf9* and *wtf33* were competed at an allelic locus in *Sp* (24). Heterozygous aneuploid or heterozygous diploid gametes, however, inherit both drivers and should be immune to both Wtf poison proteins. *Sp* (and presumably other isolates) only tolerates aneuploidy of chromosome 3, so that the fitness costs of competing drivers could be uniquely offset on chromosome 3 (22).

It is not clear why antidote-only *wtf* genes that act as suppressors of drive should specifically spread or be maintained on chromosome 3. Loci on this chromosome bear the greatest fitness cost of drivers. This is because sites on chromosome 3 are more likely to be linked in repulsion (i.e. on opposite haplotypes) to drivers that will destroy gametes that inherit them instead of the driver in heterozygous crosses. However, suppressors of drive are predicted to be favored at any unlinked locus because they increase fertility (7). It is therefore surprising that antidote-only *wtf* genes have not spread throughout the genome. We favor a model in which the frequent gene conversion amongst *wtf* genes likely leads to toggling between driving and suppressing *wtf* genes at any given locus. For example, we predict that the *wtf18* gene in FY29033 is a driver, but the *wtf18* alleles in *Sp* are suppressors of drive (Figure 2A) (11). This toggling could lead to selective maintenance of *wtf* suppressor loci on chromosome 3 due to the mechanism described above for drivers.

### Rapid evolution of *wtf* genes

We observe three mechanisms driving innovation in *wtf* gene sequences. First, as observed by Hu et al. who previously assayed *Sp* and CBS5557, we found pervasive nonallelic gene conversion affecting most *wtf* genes (24). We demonstrated that this nonallelic gene conversion was not restricted to a specific portion of the genes and included promoters. The forces driving this gene conversion will require further investigation. It is possible that the *wtf* genes inherently undergo gene conversion at a high rate due to some intrinsic property. For example, the close proximity of a subset of *wtf* loci to meiotic DSB hotspots could facilitate nonallelic recombination within the family. It is also possible that the novel *wtf* sequences generated by gene conversion are frequently advantageous. For example, novel variants could drive or suppress other drivers and thus be maintained by selection.

Second, we found that the number of units of repeat sequences within exons 3 and 6 varies greatly. Such repetitive sequences are known to be unstable and several *wtf* alleles that are otherwise very similar vary in repeat copy number. Although the function of these repeat regions is not clear, the repeats often overlap predicted transmembrane domains, and repeat number can be functionally important. For example, *Sp wtf18* antidote alleles were only able to neutralize *Sp wtf13* poison alleles that had the same number of exon 6 repeats (11). It is possible that the presence of these repeats in *wtf* genes is maintained, at least in part, due to their hypermutability. A high capacity to facilitate rapid gene diversification could be beneficial in genes involved in genetic conflicts.

The third contributor to rapid *wtf* gene evolution is positive selection in at least the *wtf7* and *wtf11* genes, which show an excess of amino acid substitutions (Figure 4). Unfortunately, extensive gene conversion limited our analyses to four genes. The *wtf7* and *wtf11* genes have no known functions and are both highly diverged from the experimentally characterized *wtf* genes and each other. The rapid evolution of these genes, however, suggests that they too are engaged in genetic conflicts. We speculate that both genes are either meiotic drivers and/or act as modifiers of meiotic drive.

### Consequences of rapid evolution

The rapid evolution of *wtf* genes has led each of the strains we assayed here to contain a unique suite of *wtf* alleles. The consequences of this *wtf* diversity on *S. pombe* fitness are profound. When nonclonal isolates of *S. pombe* mate to produce diploids, it is very likely there will be heterozygosity at one or more *wtf* loci. When these diploids undergo meiosis to generate gametes, *wtf* heterozygosity can lead to dramatic loss of fertility due to meiotic drive. This *wtf* heterozygosity is a major cause of the infertility observed in both *Sp/Sk* and *Sp*/CBS5557 heterozygous diploids and likely contributes to the generally low fertility of outcrossed (i.e. heterozygous) *S. pombe* diploids (22-24, 46, 47). Driving *wtf* genes are thus limiting the ability of *S. pombe* to enjoy all the fitness benefits of sexual reproduction, perhaps putting this species on a path to extinction.

### Lessons for the design of gene drives

The themes we describe for *wtf* gene evolution may be instructive for designing gene drives. Gene drives are engineered drive systems used to control natural populations. The general idea is that natural or artificial drivers can be used to spread traits (e.g. disease resistance) throughout a population or to eliminate a population, for example by generating extreme sex ratio imbalances (48). Analyses of natural drivers and drive suppressors, such as those of the *wtf* family, may prove useful for predicting how engineered gene drives (particularly gamete killers) may evolve if released in natural populations. For example, compact gene drives may duplicate to novel loci within a genome. This risk may be particularly high if the gene drives are integrated next to transposons or other dispersed repetitive elements.

## Materials and Methods

### Yeast strains and whole genome sequencing

The *Sp* (SZY643) and *Sk* (SZY661) strains are described in Nuckolls and Bravo Nunez et al. (23). We obtained all other strains from the National BioResource Center, Japan. We prepared genomic DNA using QIAGEN Genomic-tips (Catalog number 10262 and 10243) using the QIAGEN DNA buffer set (Catalog number 19060). We followed the kit protocol except that we extended the lyticase treatment to 18 hours and the RNase A/Proteinase K treatment to 5 hours. The Stowers Institute Molecular Biology core facility prepared the sequencing libraries using the Illumina Nextera Mate-Pair Sample Prep Kit (Cat. No. FC-132-1001). 5-8 kb fragments were selected using a BluePippin machine (Sage Science). The libraries were sequenced (150 base pair paired-end reads) on an Illumina MiSeq using the MiSeq Reagent Kit v2 (300 cycle) (Cat. No. MS-102-2002). Sequence data are available in SRA (accession no. PRJNA476416).

### Assaying *wtf* gene numbers

We used Geneious version 10.0.7 (https://www.geneious.com) for all sequence analyses, unless otherwise stated, using the ‘map to reference function’ for all short-read alignments. To find *wtf* loci in *Sk*, we identified read pairs from the mate-pair library in which one (or both) reads aligned to a library containing the 25 *Sp wtf* genes (‘medium-low sensitivity’ aligner setting) (Steps 1 and 2 in Figure 1A). For the other genomes, we also included the *Sk wtf* genes as reference sequences. From those wtf-matching read pairs, we then isolated any ‘partner’ reads that did not align to *wtf* genes by again mapping reads to our reference set of *wtf* genes (‘medium sensitivity’ setting), this time saving only the individual reads that failed to align to any *wtf* gene (Figure 1A Step 3). We then took these ‘wtf-partner’ reads and aligned them to the *Sp* reference genome (‘medium sensitivity’ setting) (Figure 1A Step 4). This generated pileups of reads flanking *wtf* loci. We inspected the pileups manually to infer the number of *wtf* genes at each locus based on the width and pattern of the pileups, as described in the text. For *Sk* and FY29033 these inferences were confirmed or corrected by assembling the *wtf* loci (see below).

### Assembling *wtf* genes

To assemble the *wtf* gene(s) at a given locus, we used flanking unique sequences as ‘bait’ to identify all read pairs in the region, and then performed individual *de novo* assemblies for each *wtf* locus separately. This approach should avoid misassemblies that can occur in whole genome assemblies at repetitive regions like *wtf* loci. In more detail, we first manually identified coordinates of the sequence pileups described above, adding ~2 kb flanking sequence (Figure 1D, orange bars under the pileups). We excluded LTR sequences and other repetitive DNA sequences from these regions and denote them ‘orange regions’. We identified all mate-pairs that align to these orange regions (‘medium low sensitivity’ setting) (Figure 1D, Step 1). We then filtered those reads so that we retained only candidate *wtf* locus reads, and not those from flanking regions. To do this, we defined two additional reference regions flanking the *wtf* locus (‘green regions’) that extend the orange region to within ~500 bp of the *wtf* locus and by ~15 kb in the other direction (Figure 1D, green bars under the pileups). We then aligned the read pairs defined in Step 1 to the green regions (‘medium sensitivity’ setting), retaining only individual reads that failed to align to the green regions; these reads represent candidate *wtf* locus reads (Figure 1D Steps 2 and 3). Finally, we assembled these candidate *wtf* reads using the Geneious ‘de novo assemble’ function (‘medium sensitivity’ setting) (Figure 1D Step 4). We obtained 1-4 contigs in most of these assemblies that we were able to stitch together manually using known *wtf* gene orientations and sequence overlaps. Gene sequences and annotations are available in GenBank (accession no. MH837193-MH837230 and MH837431-MH837459).

### DNA sequence alignments, tree construction and sequence logos

We aligned DNA sequences of the full length *wtf* genes (or of *wtf* gene segments) in Geneious using the Geneious aligner with the ‘global alignment without free end gaps’ setting. All alignments are provided as supplemental data. We then generated trees in Geneious using the PHYML plugin (version 2.2.3) with the default settings (HKY85 substitution model, set to optimize tree topology branch length and substitution rate, NNI topology search) with 100 bootstraps. For exons 3 and 6, we aligned only sequences downstream of the repetitive regions found near the beginning of those exons (Figure 3). For *wtf* family-wide gene conversion analysis, we used ran a command-line version of the GARD algorithm (using the general discrete model of site-to-site rate variation with 3 rate classes) (37). We used Weblogo3 (http://weblogo.threeplusone.com) to generate sequence logos of the repetitive regions (49).

### Analysis of selective pressures

For the initial dN/dS analyses, we first used Geneious to generate a codon alignment of the *wtf7*, *wtf11*, *wtf14* and *wtf15* genes using the ‘translation align’ function with the default settings. We then used codeml executed from PAML 4.8 to estimate dN, dS and dN/dS (runmode -2, seqtype 1, CodonFreq 0, model 0, NSsites 0, icode 0, fix_kappa 1, fix_omega 0, and omega 0.5) (43).

For the extended analyses, we mapped paired-end reads from 54 additional *S. pombe* strains to the *Sp* reference genome to generate consensus sequences of *wtf7*, *wtf11*, *wf14* and *wtf15* in the additional strains (33). The assembled sequence of these genes are available in GenBank (accession no. MH837181-MH837192 and MH837231-MH837430). We then codon-aligned a total of 57 sequences for each gene, and removed redundant sequences from each alignment using a custom script. We used the GARD algorithm (via the DataMonkey website) to screen each alignment for evidence of gene conversion (using the general discrete model of site-to-site rate variation with 3 rate classes) (37). GARD did not find evidence for gene conversion in the *wtf7*, *wtf11*, *wtf14* or *wtf15* alignments. We also used our alignments of *wtf7*, *wtf11*, *wtf14* and *wtf15* as input into the BUSTED algorithm (44), run via the DataMonkey website, to test for positive selection on a subset of sites in a subset of lineages. We also generated phylogenies from each alignment using PHYML (50) (GTR substitution model, 4 substitution rate categories, estimating the proportion of invariant sites) and used these trees and alignments as input to the codeml algorithm from the PAML package (43). We compared model 8 (M8), which allows positive selection at a subset of sites, to each of two control models, model 7 (M7) or model 8a (M8a). In each case we compared twice the difference in log-likelihoods between the two models with a chi-squared distribution with 2 (M8 vs. M7) or 1 (M8 vs. M8a) degrees of freedom to obtain a p-value. We used the F61 codon model and a starting dN/dS of 1, but also verified that our findings of positive selection in *wtf7* and *wtf11* are robust to the use of alternative parameters (codon model F3×4, and starting omega of 0.4 or 3 for M7 and M8 - starting omega cannot be varied for M8a).

## Acknowledgements

The authors thank Blake Billmyre and other members of the Zanders laboratory for helpful discussions and for comments on the manuscript. Gene sequences and annotations are available in GenBank (accession no. MH837181-MH837459). This work was supported by the Stowers Institute for Medical Research and the National Institutes of Health (NIH) award number R00GM114436 (SEZ). JMY is supported by NIH award number R01GM074108 to Harmit S. Malik.

## Competing interests

SEZ: Inventor on patent application based on *wtf* killers. Patent application serial 62/491,107. MTE and JMY declare that no competing interests exist.

## Figure Legends

**Table 1: Meiotic double strand break hotspots are enriched near *wtf* genes.**

**Supplemental Table 1:** The names previously given to each CBS5557 *wtf* gene by Hu et al. are shown on the left and the names we use in this work are shown on the right.

**Supplemental Figure 1**: Representative long RNA sequence reads (dark grey) from Kuang et al. are shown aligned to *Sp wtf* genes (30). The solid lines represent UTRs, boxes are exons and the thin lines are introns (i.e. they were not found in the reads). The blue gene annotations are from PomBase. The light grey annotations are based on the long read RNA sequencing reads. The light grey annotations were used for the coding sequences in this work. Predicted pseudogenes have an ^*^ after the gene name. On the annotations, the light grey annotations used in this work, the ‘^*^’s indicate in frame stop codons and the ‘/’s indicate frameshift mutations.

**Supplemental Figure 2:** Insert sizes of mate-pair libraries. Most pairs of reads map as expected for mate-pair fragments (i.e. the 3’ end of the reads point away from each other in the genome) with insert size ~ 6-10 kb. These were the reads that were useful in identifying and assembling *wtf* loci. However, the mate-pair library prep is not 100% efficient, and also generates a subset of reads with inserts of < 1 kb that typically map as regular paired-end reads (i.e. the 3’ end of the reads point towards each other in the genome); these reads were generally discarded by the selective steps of our sequence analysis pipelines (Figure 1A and 1D).

**Supplemental Figure 3:** The width of the sequence pileups used to identify loci containing *wtf* genes and to predict the number of *wtf* genes at each locus. The width of both the 5’ and 3’ pileups is shown for loci with one, two, or three verified *wtf* genes from *Sp* (blue), *Sk* (red), and FY29033 (green). In general, the widest pileup at a given locus was used to predict the number of *wtf* genes when the pileups were asymmetric (e.g. in the loci with three *wtf* genes).

**Supplemental Figure 4:** DNA sequence read pileups flanking *wtf* loci for atypical loci, showing representative of the patterns not shown in Figure 1B. In general, the atypical pileup patterns were caused by Tf transposon insertions in the sequenced strain that were not present in the *Sp* reference genome to which the reads were aligned. These transposon insertions make the actual genome from which the reads were derived different from the reference genome and unique sequences (not transposons) are needed next to a *wtf* locus to form a pileup. The Tf insertions were discovered during assembly of the loci. Black arrows indicate the locations of *wtf* genes and green arrows represent Tf transposons.

**Supplemental Figure 5:** Sequence coverage of the *wtf23* region in FY29030 and the *wtf33* region in FY28989. We infer from the roughly doubled coverage of those regions that they are duplicated in those strains. In addition, high sequence identity across the duplicated regions within each strain are consistent with recent duplications and very little divergence between the two copies.

**Supplemental Figure 6:** The *wtf* gene names are shown mapped onto the karyotype of *Sp*, although not all the strains share this karyotype. Genes on the Watson strand are shown above each chromosome, whereas genes on the Crick strand are shown below chromosomes. Experimentally confirmed drivers and genes we predict to be intact drivers are shown in purple text. Predicted pseudogenes are indicated with an asterisk.

**Supplemental Figure 7:** Maximum likelihood tree generated by PhyML (executed in Geneious) including full-length ORF sequences of all *wtf* genes from *Sp*, *Sk*, FY29033 and CBS5557 (alignment length 1,465 bp). The tree is unrooted, but is shown with arbitrary rooting in (A) to facilitate reading the branch labels. Predicted pseudogenes are indicated with an ^*^. Nodes with ≥95& bootstrap support are indicated with red circles. The same tree is shown unrooted in (B). The *wtf7* (dark blue), *wtf11* (pink), *wtf14* (brown) and *wtf15* (light blue) clades are each highlighted. The clade highlighted in green was discussed in the text as an example. The scale bar indicates nucleotide substitutions per site.

**Supplemental Figure 8:** GARD analysis of all *wtf* ORF sequences from *Sp*, *Sk*, FY29033 and CBS5557. This analysis found that a hypothesis allowing multiple trees for different segments of the alignment is >100 times more likely than a hypothesis allowing only a single tree, supporting that recombination operates within *wtf* genes. The analysis identified two likely breakpoints corresponding to positions 615 and 1047 in the alignment, yielding three segments as depicted by the colored rectangles at the top of the figure. Both breakpoints have strong statistical support (^***^; p<0.01). The trees generated for each segment (below) are distinct.

**Supplemental figure 9:** Maximum likelihood tree for exon 1 of the *wtf* genes (alignment length 150 bp). See legend to Supplemental Figure 7 for details. The grey shaded box corresponds to the black color coded exon in the cartoons in Figure 2A.

**Supplemental figure 10:** Maximum likelihood tree for exon 2 of the *wtf* genes (alignment length 381 bp). See legend to Supplemental Figure 7 for details. The grey shaded box corresponds to the black color coded exon in the cartoons in Figure 2A.

**Supplemental figure 11:** Maximum likelihood tree for exon 3, excluding repeats, of the *wtf* genes (alignment length 203 bp). See legend to Supplemental Figure 7 for details. The grey shaded box corresponds to the black color coded exon in the cartoons in Figure 2A.

**Supplemental figure 12:** Maximum likelihood tree for exon 4 of the 6-exon *wtf* genes (alignment length 192 bp). See legend to Supplemental Figure 7 for details. The grey shaded box corresponds to the black color coded exon in the cartoons in Figure 2A.

**Supplemental figure 13:** Maximum likelihood tree for exon 5 from the 6-exon *wtf* genes and the homologous exon 4 of the 5-exon *wtf* genes (alignment length 65 bp). See legend to Supplemental Figure 7 for details. The grey shaded box corresponds to the black color coded exon in the cartoons in Figure 2A.

**Supplemental figure 14:** Maximum likelihood tree for exon 6, excluding repeats, from the 6-exon *wtf* genes and the homologous exon 5 of the 5-exon *wtf* genes (alignment length 69 bp). See legend to Supplemental Figure 7 for details. The grey shaded box corresponds to the black color coded exon in the cartoons in Figure 2A.

**Supplemental figure 15:** Maximum likelihood tree of the region upstream of *wtf* genes (alignment length 303 bp). See legend to Supplemental Figure 7 for details. The grey shaded box corresponds to the black color coded region in the cartoons in Figure 2A.

**Supplemental figure 16:** Maximum likelihood tree for intron 1 of the *wtf* genes (alignment length 280 bp). See legend to Supplemental Figure 7 for details. The grey shaded box corresponds to the green color coded intron in the cartoons in Figure 2A.

**Supplemental Figure 17:** A codon alignment of the *wtf7* orthologs from *Sp*, *Sk*, FY29033 and CBS5557. Purple boxes represent exons; all DNA and amino acid sequence variants are highlighted.

**Supplemental Figure 18:** A codon alignment of the *wtf11* orthologs from *Sp*, *Sk*, FY29033 and CBS5557 is shown. Purple boxes represent exons; all DNA and amino acid sequence variants are highlighted.

**Supplemental Figure 19:** A codon alignment of the *wf14* orthologs from *Sp*, *Sk*, FY29033 and CBS5557 is shown. Purple boxes represent exons; all DNA and amino acid sequence variants are highlighted.

**Supplemental Figure 20:** A codon alignment of the *wtf15* orthologs from *Sp*, *Sk*, FY29033 and CBS5557 is shown. Purple boxes represent exons; all DNA and amino acid sequence variants are highlighted.

